# Brain mapping across 16 autism mouse models reveals a spectrum of functional connectivity subtypes

**DOI:** 10.1101/2020.10.15.340588

**Authors:** V. Zerbi, M. Pagani, M. Markicevic, M. Matteoli, D. Pozzi, M. Fagiolini, Y. Bozzi, A. Galbusera, ML Scattoni, G. Provenzano, A. Banerjee, F. Helmchen, M. Albert Basson, J. Ellegood, J. P. Lerch, M. Rudin, A. Gozzi, N. Wenderoth

## Abstract

Autism Spectrum Disorder (ASD) is characterized by substantial, yet highly heterogeneous abnormalities in functional brain connectivity. However, the origin and significance of this phenomenon remain unclear. To unravel ASD connectopathy and relate it to underlying etiological heterogeneity, we carried out a bi-center cross-etiological investigation of fMRI-based connectivity in the mouse, in which specific ASD-relevant mutations can be isolated and modelled minimizing environmental contributions. By performing brain-wide connectivity mapping across 16 mouse mutants, we show that different ASD-associated etiologies cause a broad spectrum of connectional abnormalities in which diverse, often diverging, connectivity signatures are recognizable. Despite this heterogeneity, the identified connectivity alterations could be classified into four subtypes characterized by discrete signatures of network dysfunction. Our findings show that etiological variability is a key determinant of connectivity heterogeneity in ASD, hence reconciling conflicting findings in clinical populations. The identification of etiologically-relevant connectivity subtypes could improve diagnostic label accuracy in the non-syndromic ASD population and paves the way for personalized treatment approaches.

## Introduction

Autism spectrum disorder (ASD) is a neurodevelopmental condition marked by social, communication and behavioral challenges often accompanied by additional co-morbidities that together negatively impact the quality of life of affected individuals and their families. The high heterogeneity of clinical presentation and underlying pathophysiology pose a substantial challenge for early diagnosis and effective treatments ^1,2^. Among the multiple and diverse etiological factors associated with ASD ^3^, genetic alterations appear to be by far the largest contributors to ASD risk ^4^. Several studies have revealed that genetic variants associated with ASD cause cellular alterations linked to abnormal neuronal circuit wiring and function, leading to aberrant developmental trajectories (reviewed by ^5^). Hence, circuit and network dysfunctions are thought to directly underlie onset and severity of ASD symptoms.

Within this framework, abnormalities in the coordinated functional interactions of brain networks, or brain connectivity, might therefore represent a defining hallmark of ASD. This theoretical view was first inferred from fMRI measurements of cortical activation collected during different cognitive tasks^6^. Since then, a growing number of studies in idiopathic ^7,8^ as well as syndromic forms of ASD ^9–12^ has suggested that aberrant connectivity in ASD could be detected by resting-state fMRI (rsfMRI). However, whilst a recent aggregate analysis has revealed a putatively reproducible mosaic pattern of atypical connectivity in ASD^13^, the heterogeneity in rsfMRI connectivity findings across ASD cohorts is considerable, and unlikely to reflect technical or procedural discrepancies in imaging acquisitions and analysis^2,12,14^. Hence, one outstanding question in the field is whether ASD can be associated with a univocal, diagnosis-specific signature of dysfunctional brain connectivity that is common to the whole spectrum, or whether clinical heterogeneity is the sum of distinct and separable signatures of network dysfunction.

rsfMRI connectivity studies in etiologically homogeneous ASD populations have started to link disease-causing genetic alterations to specific functional connectivity aberrancies, highlighting putatively distinguishable, etiology-specific connectivity changes. For example, subpopulations harboring well-characterized genetic mutations such as chromosome 16p^11^.2 deletion 11, neurofibromatosis type 1 mutations (Nf1) ^15^ or Fragile-X (Fmr1) syndrome ^9,16^ are characterized by non-overlapping patterns of dysconnectivity. Similarly, mTOR-related synaptic surplus has been recently linked to a specific cortico-striatal hyper-connectivity signature^17^. These observations suggest an interpretative framework in which ASD-relevant etiological and genetic risk factors could lead to distinct network signatures of brain dysfunction, thus explaining heterogeneous connectivity alterations observed in clinical cohorts. Crucially, this model would also explain why so far, no unequivocal ASD-specific signature of abnormal connectivity has been identified.

To test this hypothesis, we introduce the Autism Mouse Connectome (AMC) study, a bi-center initiative dedicated to collecting and analyzing rsfMRI in multiple ASD-relevant mouse mutants under well-controlled and highly reproducible experimental conditions^18–20^. Our work leverages recent advances in cross-species rsfMRI imaging, revealing that rsfMRI is exquisitely sensitive to network alterations caused by ASD-related etiologies, reconstituting patterns of network dysfunction in corresponding human ASD cohorts ^11,16 21–26 17,21–26^. By comparing connectivity alterations caused by 16 ASD-related genetic and etiological factors, we found that ASD etiologies cause a wide spectrum of connectivity abnormalities that can be clustered into a discrete set of prevailing subtypes. Our results underscore a pivotal contribution of etiological variability to connectivity heterogeneity in ASD and reveal a set of prominent cross-etiological network dysfunction modes of high translational relevance for ASD.

## Results

### Animal models

The AMCB collection includes scans from published literature that have been retrospectively aggregated and additional unpublished work (see Table 1). In total, it contains resting-state fMRI scans of 350 mice from 16 distinct cohorts. Ten cohorts were collected at ETH Zürich (Switzerland) and six cohorts were collected at IIT Rovereto (Italy). Data from two independent cohorts of CNTNAP2 were collected at both sites. All mouse mutants fulfilled the following criteria: 1) the genetic modification resembles/relates to a genetic alteration found in individuals with ASD as listed in the SFARI gene database (https://gene.sfari.org/autdb/); 2) the mouse strain has been shown to mimic at least one of the core behavioural phenotypes of ASD; 3) mice are inbred with C57BL/6J or /6N mice as control strain to reduce genetic heterogeneity across models; 4) each cohort included mice with the genetic alteration and wild-type control littermates. In addition, we included: (i) a model for environmental ASD risk factor, i.e. maternal exposure to interleukin-6 (IL-6); the maternal injection of IL-6 at later stages of pregnancy (E12,5-E15) is a well-established model of Maternal Immune Activation (MIA), and the relative offspring show major behavioural defects which include alterations in sensory-motor gating, attention and sociability, all clear hallmarks of ASD ^27,28^; (ii) a model for TREM2 deficiency; TREM2KO mice are characterized by defects in microglia-dependent synaptic pruning activity which results in excessive glutamatergic synaptic connections accompanied by autistic-like behavioural phenotype ^29^ (iii) a model of congenital agenesis of the corpus callosum, a neuroanatomical trait associated with increased prevalence of ASD ^30^; inbred BTBR mice are often employed as a rodent model for “idiopathic” ASD and exhibit profound behavioural and neurofunctional alterations of high translational relevance ^31–33^.

**Table 1.** Available datasets.

| ASD mouse model | Control Strain | Sex | Age (weeks) | Behavioral traits * | Construct validity † | References | Scan Site |
| --- | --- | --- | --- | --- | --- | --- | --- |
| 16p11.2 (df/+) | 129/C57BL6 J | males | 18-20 | 3,5 | c | <sup>11</sup> | IIT |
| BTBR T+tf/J | C57BL/6J | males | 26 | 1,2,3,4,5 | d | <sup>34</sup> | IIT |
| CDKL5 KO | C57BL/6J | males | 23-25 | 3,5 | b,f | - | ETH |
| CDKL5 Het (+/-) | C57BL/6J | females | 17-19 | 3,5 | b,f | - | ETH |
| CHD8 | C57BL/6J | mixed | 15-18 | 3,4,5 | a | <sup>26</sup> | IIT |
| CNTNAP2_ETH (-/-) | C57BL/6J | mixed | 14-16 | 1,4,5 | a,f | <sup>16</sup> | ETH |
| CNTNAP2_IIT (-/-) | C57BL/6J | males | 13-14 | 1,4,5 | a,f | <sup>23</sup> | IIT |
| En2 (-/-) | C57BL/6J | mixed | 12-16 | 1,4,5 | a,e,d | <sup>24</sup> | ETH |
| FMR1.1 (-/y) | C57BL/6J | males | 14-16 | 1,2,3,4 | b,f | <sup>16</sup> | ETH |
| FMR1.2 (-/y) | C57BL/6J | males | 13-15 | 1,2,3,4 | b,f | <sup>35</sup> | ETH |
| IL6 | C57BL/6J | mixed | 14 | 1,4 | d | - | ETH |
| Mecp2 Het (+/-) | C57BL/6J | females | 12-14 | 1,4,5 | b,f | - | ETH |
| SGSH (-/-) | C57BL/6N | mixed | 20-22 | 5 | b | - | ETH |
| SHANK3b ex4-9 | C57BL/6J | males | 19-21 | 1,2,3,4,5 | a,f | <sup>25</sup> | IIT |
| Syn2 | C57BL/6J | males | 25-31 |  | f | <sup>36</sup> | IIT |
| TREM2 (-/-) | C57BL/6J | mixed | 12-15 | 3,4 | d | <sup>29</sup> | ETH |

\* *Behavioral traits related to ASD reported by others in selected models: (1) social behavior deficits; (2) reduced vocalizations; (3) repetitive behavior; (4) cognitive behavior; (5) motor performance. † Construct validity includes: (a) Genetic association; (b) Related syndrome; (c) Autism copy number variants; (d) Autistic-like behavior; (e) Serotonin-related phenotype; (f) Synapse related*

### Between-cohorts and between-sites connectome consistency in control animals

rsfMRI data from all cohorts were commonly preprocessed using a standard minimal-preprocessing pipeline and normalized to the Allen Mouse Brain Common Coordinate Framework (CCFv3) (Figure 1A). Our first goal was to test the assumption that scans from wild-type (WT) control animals in each of the cohorts would show comparable functional connectome characteristics. A total of 112 datasets from WT animals were selected and used to compute an averaged rsfMRI mouse connectome (Figure 1B). The similarity between the connectome representation of each individual and the group average of all WTs -expressed by Spearman’s ranking coefficient (Rho) -is used to quantify consistency of the data (larger Rho refers to larger similarity between WTs). This parameter was evaluated for a range of sparsity levels of the functional connectome which varied from keeping only the top 1% of the strongest and positively correlated edges to keeping up to 50%. Maximum similarity between datasets was obtained at 4% sparsity (545 edges) (Figure 1C). This indicates that the functional connectome in the mouse is maximally similar between all data sets when considering the 4% strongest and positively correlated edges. This level of sparsity was applied to all subsequent analyses. Similarity indexes between the individuals of each cohort and the group average of all WTs are shown in Figure 1D. The results show a marked resemblance across animals (mean Spearman’s rho: 0.699 ± 0.06) and low variability between cohorts.

**Figure 1.**
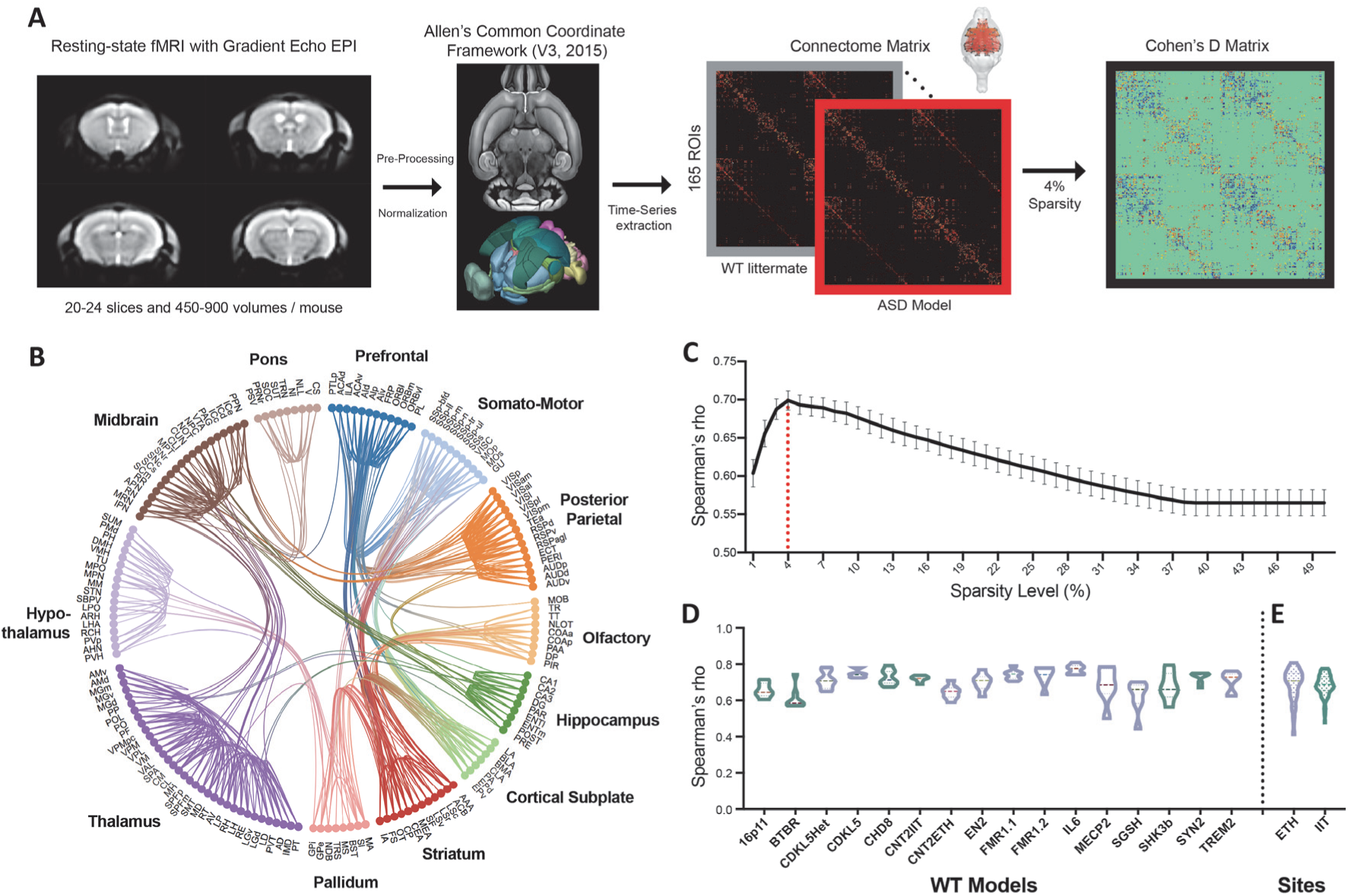
WT datasets show comparable connectome representation across recording sites. (A) Schematic of the experimental pre-processing and processing pipeline. (B) Circos plot of the mouse brain connectome from 112 rsfMRI wildtype datasets, 4% sparsity threshold, highlights a complex network of distributed short-and long-range connections (number of edges = 545). (C) Quantification of similarity across all WT mice at different sparsity levels (1-50%). Maximum averaged Spearman’s rho was found at 4% sparsity. (D) Distributions of ranked similarity in connectome representation (Spearman’s rho) between wildtype of each cohort and the group average (E) Same data as in D but grouped within each of the two measurement sites (ETH, IIT). Both nonparametric testing and Bayesian repeated-measures ANOVAs show that datasets from WT mice exhibit comparable connectome representation independently from recording site. Wilcoxon matched-pairs signed rank test, p-value = 0.2749. Bayesian factor BF-H0 (null) =7.57.

Next, we tested whether the data across the two sites exhibit significant differences, for example, due to differences in environmental conditions in which mice were raised, experimental protocols or data acquisition hardware. We collected an equal number of wildtype animals for each site (n=75) randomly distributed across cohorts and measured their similarity to a common template with Spearman’s Rho. Importantly, we found no significant differences in mean similarity between data collected at the two sites (Wilcoxon matched-pairs signed rank test, *p*-value = 0.2749, Figure 1E). In addition, Bayesian repeated-measures ANOVAs revealed a Bayes Factors in favor of the null hypothesis (BF-H0, i.e. data from the two sites are *not* different) of 7.57 versus a BF that rejects the null hypothesis of 0.13 (BF-H1, i.e. data from the two sites are different). Overall, these data clearly indicate that the connectome representation is comparable in WT animals irrespective of which site performed the rs-fMRI measurements (ETH and IIT).

### Low-dimensional representation of connectivity changes in individual animals reveals a wide spectrum of alterations

Autism is commonly defined as a "spectrum" of neurodevelopmental disorders, owing to the wide range of etiologies and pathophysiological factors that has been associated with this condition. It is however unclear whether abnormal functional connectivity faithfully mirrors the ASD etiological variability or converges onto a single common signature of circuit dysfunction. To disambiguate these two opposing possibilities, we implemented a dimensionality reduction algorithm and mapped individual mouse connectivity data into the same coordinate system. After determining connectivity deviations in the ASD mutants relative to their own WT controls, connectivity abnormalities of each animal were projected onto a low-dimensional space using an unsupervised manifold learning technique (Uniform Manifold Approximation and Projection, UMAP), which preserves the global data structure and the local neighbor relations better than other existing methods such as Principal Components or t-distributed Stochastic Neighbor Embedding (t-SNE) ^37^. Point-to-point distances in UMAP plots were then used to interpret the continuity of the data subsets, and identify similarities in ‘connectopathy’ between individual animals. Notably, the resulting map revealed a continuous spectrum of connectivity abnormalities across which individual etiological clusters could be located and identified (2D representation, Figure 2). The analysis of the Euclidian distance between points in the low-dimensional embedded space (corresponding to the (dys)connectivity profiles of individual animals) revealed that distances were shorter between individuals of the *same* ASD cohort (reflecting more similar connectivity alterations) than between individuals of *different* ASD cohorts (for 2D embedding, Euclidean distance within-groups = 1.5 ± 0.56; between groups = 2.49 ± 0.45, T-test p = 2.55e^−8^, Supplementary Figure 1). This feature was preserved when using a higher number of dimensions in the embedding space (three to ten dimensions, Supplementary Figure 1). Importantly, the overall distribution of subjects across this low-dimensional space defined a cross-etiological continuum within which some ASD etiologies appeared to be clustered and distinguishable (e.g. FMR1, IL-6, CDKL5, BTBR) while others were widespread and partly overlapping. Taken together, these results suggest that i) the underlying etiology determines the similarity in connectopathy between individual animals and their position in the embedded space; ii) the resulting distribution of connectivity profiles defines a continuous landscape of connectivity alterations, arguing against the idea of a common pattern of connectivity abnormalities across all the probed models. Importantly, dimensionality-reduction method such as UMAP supports an inverse transform that can approximate how a new sample – or new animal model -would connect to a specific position in the embedding space. This feature is particularly interesting for future research that aims to compare new genetic models within the two-dimensional reference framework of this first AMC study.

**Figure 2.**
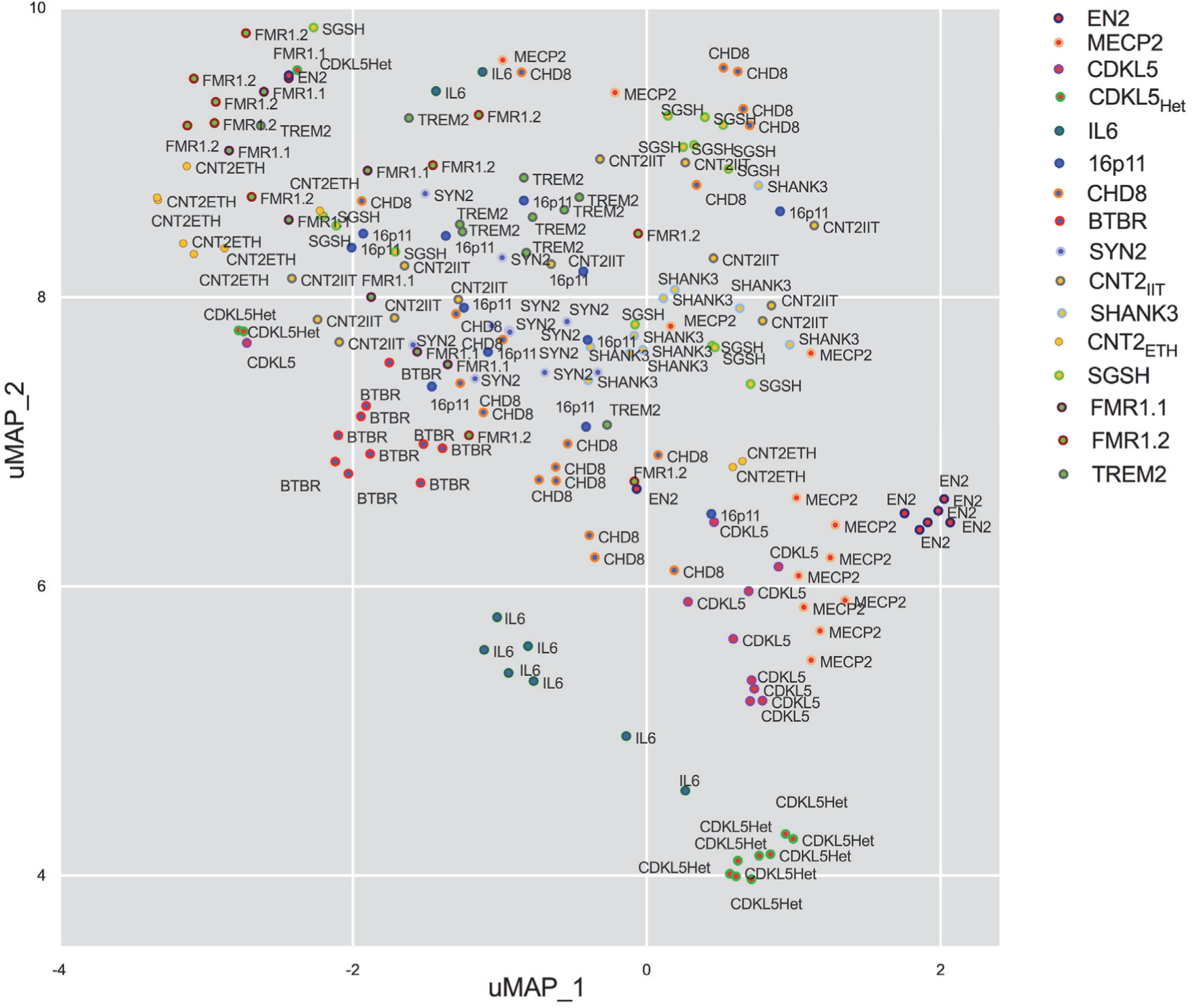
ASD-related etiologies define a continuous connectivity landscape. Uniform manifold approximation and projection (UMAP) 2-dimensional embedding of the connectome data from 176 individual ASD-related model animals of the AMC dataset. Individual data are Z-scored and normalized to the average cohort’s WT control population. The color of the elements represents the model’s cohort. uMAP parameters: n_neighbours = 10, min_dist = 0.1.

### ASD-specific connectivity signatures can be segregated into cross-etiological clusters

In the previous analysis, we have shown that different autism-associated genetic etiologies define a pseudo-continuous landscape of connectivity alterations. However, one outstanding question is how well a "categorical" diagnostic approach, for example based on quantitative and robust clustering methods, can effectively model this complexity and define common connectivity signatures across mutants 38. To answer this, we first assessed the deviations in connectivity strength for each of the edges between the mutant mice and their control littermates by their effect size, i.e. Cohen’s *d*. The Cohen’s d was then entered into a matrix (545 edges x 16 cohorts) representing connectivity alterations for each of the 16 models (Figure 3A). In keeping with the results of our low dimensional mapping, a visual inspection of the obtained matrix revealed the presence of different connectivity profiles, entailing spatially distributed rather than focal patterns of abnormal connectivity. Some of the abnormal patterns exhibited opposing features, such as over-and under-connectivity within the same pair of nodes (see top left and bottom left section of matrix in Figure 3A). The resulting matrix was fed into an unsupervised Gaussian Mixture Model (GMM) clustering algorithm to determine the general structure of the data and to establish putative similarities between the different mouse cohorts. In order to verify the reproducibility and stability of these clusters, we measured the proportion of time that two cohorts were clustered together using a bootstrap procedure (Figure 3B). Therefore, we randomly selected 80% of the data within each cohort, calculated the Cohen’s d for all edges, and applied the clustering algorithm (1000 times). We also determined a null distribution by repeating this procedure on data that were randomly shuffled between mutants and WT (also 1000 times, Figure 3C). To determine the optimal number of clusters, we computed the *silhouette* score for a number of possible cluster solutions (ranging from 2 to 16) using GMM. The silhouette score considers the distance between one and all other cohorts *within* the same cluster (cohesion), and the distance between one and all other cohorts in the *next nearest* cluster (separation). High *silhouette* values reflect high cohesion and high separation indicate that cohorts are well matched to their own cluster and poorly matched to neighboring clusters. Since the GMM procedure is not deterministic, we ran 1000 fits for each number of clusters. The difference in silhouette scores between the bootstrapped and the null model was used to determine the best cluster solution(s). The bar plot in Figure 3D shows a high difference score for two, three and four-cluster solutions. In order to preserve most of the complexity in the data, we kept the four-cluster solution for the following analyses. Next, we calculated the connection strength probability between each pair of cohorts (i.e. how often they have been assigned to the same cluster during bootstrapping; Figure 3B, E). When grouping the cohorts into four clusters, the cluster with the strongest within-cohort connections is formed by TREM2 and both Fmr1 cohorts, which were grouped together on average 74% of the time. This was followed by a cluster that includes CNTNAP2, Shank3b, and SGSH (on average grouped 73% of the time). The two CNTNAP2 models scanned at different institutes were clustered together 75% of the time. The other two clusters include (i) CDKL5 KO, CDKL5 Het, Mecp2 Het and EN2 (59%, with CDKL5 KO and CDKL5 Het being clustered together 67% of the time) and (ii) BTBR, IL6, 16p11 and CHD8 (on average grouped 51% of the time). The hierarchical representation of the clustering connections between cohorts is shown in Figure 3E. UMAP representation of individual animals color-coded by their respective cluster is shown in Supplementary Figure 2. Taken together, these findings suggest that ASD-specific connectivity signatures can be segregated into distinguishable cross-etiological clusters.

**Figure 3.**
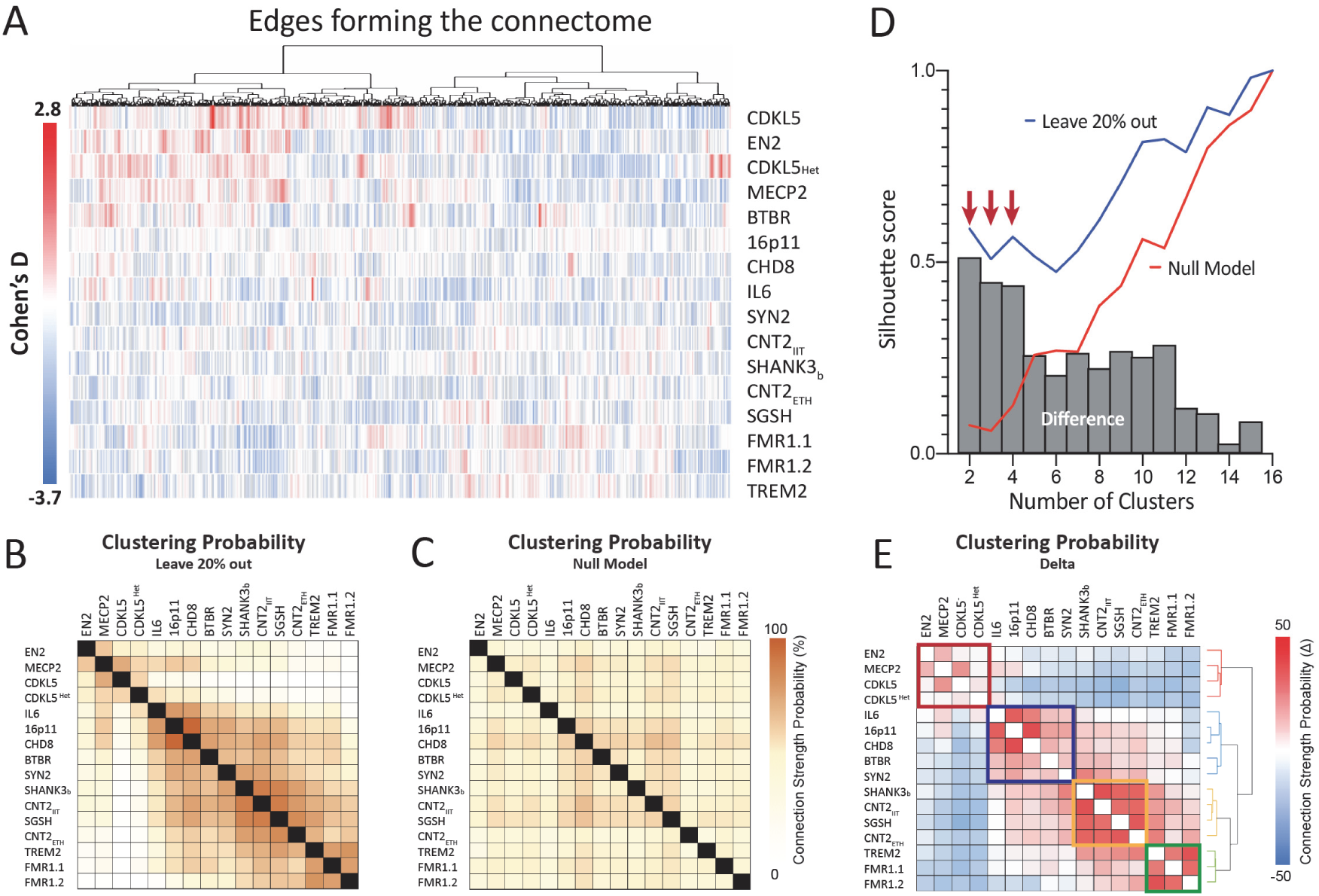
Functional connectivity signatures can be grouped into four cross-etiological clusters. (A) Functional connectivity aberrances in the 16 ASD mouse cohorts. The heatmap displays the effect size (Cohen’s D) differences in connectivity strength between the 16 different mouse models and their specific WT controls for each of the 545 different edges across the connectome. Red represents over-connectivity compared to control and blue represents under-connectivity. The dendrogram on the x-axis represents the correlation between edges. (B) Gaussian Mixture Model revealed similarities across mouse cohorts. Clustering probability (%) is measured based on the proportion of time that two cohorts belong within the same cluster over the 1000 bootstrapped samples using the leave-20%-out criteria. (C) Clustering probability of the null model generated by randomly assigning knockout and wildtype labels in each cohort 1000 times. (D) Silhouette score measured mean intra-cluster distance and the mean nearest-cluster distance in both the real and null distributions for different cluster solutions (n=2, 3, … 16). High silhouette score differences are found in the 2,3 and 4 cluster solutions. (E) The hierarchical clustering using 4-cluster solution segregated the models into specific groups depending on their connectivity similarity.

### Common connectome deviations across etiologies define distinguishable ASD connectivity subtypes

Our results so far indicate that mutations in different ASD-related genes are associated with distinctive connectivity aberrances. Some of these ASD mouse models, however, exhibited similar profiles of abnormal connectivity that can be grouped and categorized into a small number of cross-etiological subtypes. To further pinpoint the spatial location of these connectivity alterations within each subtype, we used a general linear model (GLM) with nonparametric permutation testing to evaluate for connectivity abnormalities when all mice within one cluster were pooled. The results are shown in their anatomic structure (node-level comparison, Figure 4A). The original structure (edge-edge comparison) and their macro-area (parent-level comparison) are shown in Supplementary Figure 3.

**Figure 4.**
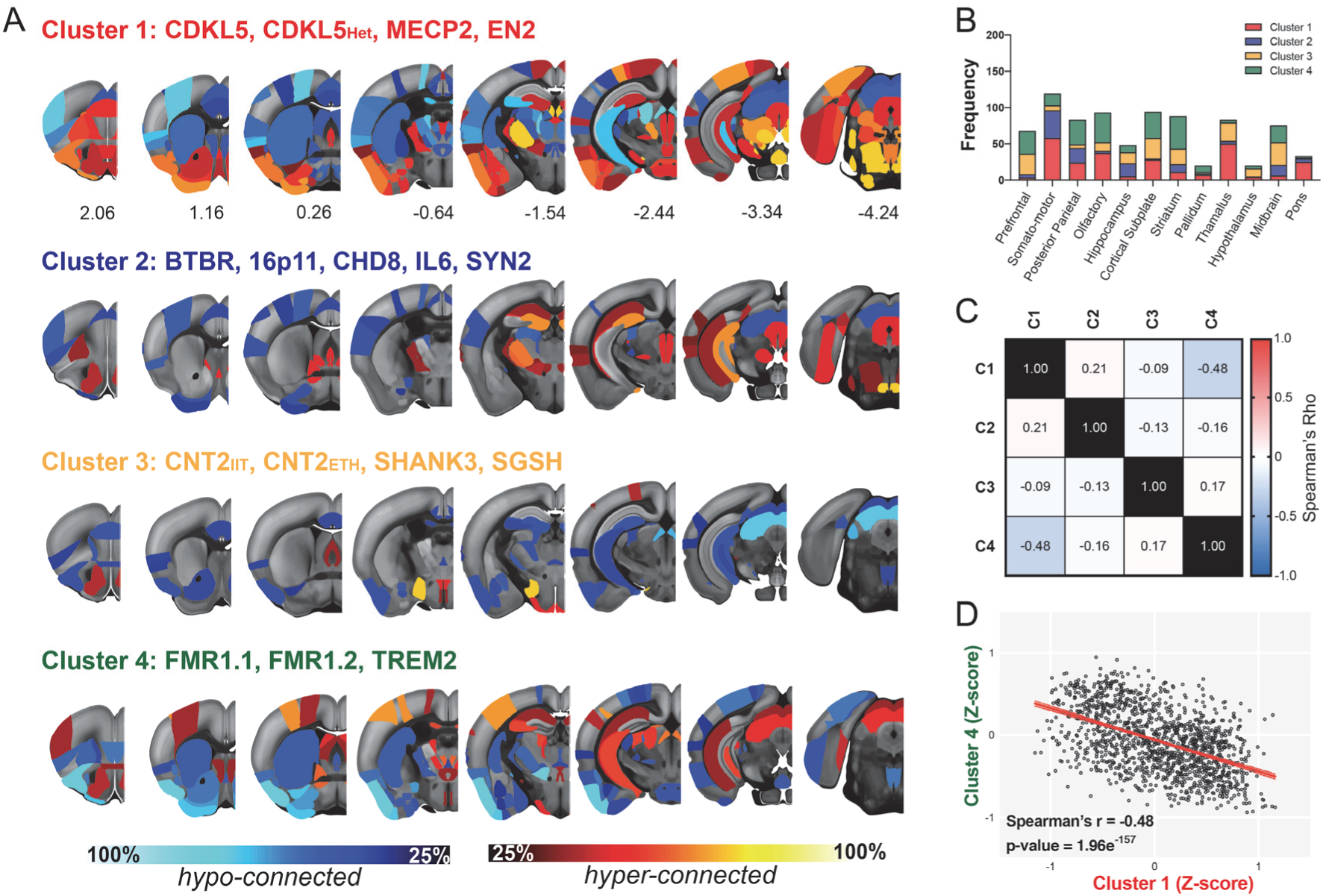
Anatomical representation of connectivity deficits in the four subtypes. (A) Rendering of regional connectivity deficits in the four clusters at the node level, revealing a heterogeneous set of brain areas with prominent over-and under-connectivity. Data are visualized in Allen Mouse reference space. (B) Number of connections (displayed as stacked frequencies) that exhibited abnormalities at the parent level. (C) Correlation matrix between all clusters, considering all 545 edges. (D) A significant negative correlation was found between Cluster 1 and Cluster 4. Spearman’s rho = −0.48, p-value = 1.96e^−157^.

As expected, the distribution of significant over-and under-connected edges (p-value < 0.05) varied across the four clusters, defining four distinct connectivity signatures (Figure 4A). The first cluster was characterized by under-connectivity in insula, somato-motor cortices, anterior cingulate, caudoputamen, hippocampus, colliculus and increased connectivity between areas in prefrontal, orbitofrontal, piriform, visual cortices, amygdala, ventral posterolateral thalamus and pontine nuclei. The second cluster showed under-connectivity between cortico-striatal areas and inferior colliculus but increased connectivity between ventral orbital, lateral septal nuclei cortex and hippocampus. The third cluster displayed under-connectivity of anterior cingulate, insula, hippocampus and thalamic VPM and only a moderate over-connectivity in the accumbens shell and hypothalamus. The fourth cluster exhibited under-connectivity between piriform and olfactory-related areas, striatum and thalamus (polymodal-associated areas) and over-connectivity in hippocampus and hypothalamus.

Across all clusters, the regions showing higher vulnerability to abnormal connectivity (independent of directionality, i.e. over-or under-connectivity) were the somatomotor regions, olfactory and cortical subplate. Pallidum, hypothalamus and pons were the least affected areas (Figure 4B). Interestingly, some of the observed connectivity features across clusters appeared to exhibit opposing configurations. Such effect was apparent in Clusters 1 and 4, where diverging connectivity alterations in cortical and hippocampal areas were observed. This notion was quantitatively corroborated by formal testing of the independence of the clusters, in which we compared the connectivity profiles in all four clusters using ranked statistics (Figure 4C). This analysis revealed no correlation among the cluster, with the predicted exception of a strong negative correlation between edges in clusters 1 and 4 (Spearman’s rho=-0.48, p=1.96e^−157^), i.e. edges that were over-connected in cluster 4 were under-connected in cluster 1 (and vice versa). This data suggests that these etiologies affect the same networks, but connectivity deviates in opposite directions (Figure 4D).

The previous analysis described which anatomical networks exhibited connectivity abnormalities in either of the four clusters. Next, we sought to identify whether we could isolate aberrant connectivity patterns that are *unique* for a given cluster, hence defining which anatomical connections are preferentially affected in each subtype. To this end, we grouped the animals in each cluster and compared it to the other models using nonparametric permutation tests. In line with the results of our regional mapping (Figure 4A), this analysis confirmed the presence of a cluster-specific functional connectivity alterations (Supplementary Figure 4).

Finally, we probed the presence of a connectivity signature that could emerge at the population level, across all the etiologies tested in our study. This was done by testing all genetic models against all wild-type littermates. Consistent with the large heterogeneity in the (dys)connectivity signatures mapped, this analysis revealed only a small number of connections (n=24, 4.4% of the selected edges) that survived correction for multiple comparisons (Supplementary Figure 5). Under-connectivity was observed between somatomotor areas, anterior insula, caudoputamen, fundus of striatum, endopiriform nucleus and claustrum, including a set of substrates that is not representative of the wide area of substrates identified in our cluster analysis. these results seem to indicate that there are only very few connectivity deficits that are common among the models, arguing once again for the presence of a common brain signature of dysfunction that is characteristic of the autism spectrum.

## Discussion

The absence of reliable and specific cross-etiological molecular or genetic biomarkers for ASD ^1,39^ have prompted research into the use of functional neuroimaging readouts as possible point of convergence for a diagnostic or prognostic characterization of ASD. However, until now connectivity studies in ASD patients have revealed highly heterogeneous and inconsistent results, which has fueled discussion regarding the clinical utility of imaging markers in ASD^40^. Here, we capitalized on recent advances in rodent rsfMRI^19,20,41–44^ to measure connectivity across 16 different ASD mouse models. We aimed to coarsely approximate the heterogeneity and etiological complexity of ASD, with the advantage of having a tight control of environmental factors, *a priori* information on the nature of these etiologies, and a well-matched reference population for connectivity mapping (i.e. WT littermate mice), hence controlling for major confounding factors in clinical neuroimaging research.

Our approach resulted in three important findings that bear high translational relevance for ASD research. First, we found that all models are characterized by significant alterations in brain functional connectivity entailing multiple, spatially distributed patterns of abnormal connectivity, rather than focal atypicalities. This finding is consistent with human literature^2^, and suggests that altered large-scale inter-areal communication is a hallmark endophenotype in ASD across etiologies, substantiating prior conceptualizations of ASD as a brain “connectopathy”^45,46^. Second, our data exhibited a broad spectrum of connectivity aberrancies even when mapped onto a low-dimensional landscape, indicating the *absence* of a prominent consensus pattern of abnormal connectivity across etiologies. This observation is of great importance in light of the ongoing debate as to the origin and significance of connectivity changes described in rs-fMRI of ASD individuals. Together with the results of analogous cross-etiological clustering of brain structure in rodent ASD models ^47^, our data argue against the existence of a reliable and specific autism-specific brain signature, and suggest that the substantial heterogeneity underling the neurobiology is a plausible key driver for the multiple inconsistent connectivity findings in clinical ASD. Finally, a third central conclusion of our analysis is that different ASD etiologies can be grouped into a small number of “families” or "clusters" which exhibit common atypicality in specific brain connections, defining a putative set of network dysfunction ASD subtypes ^2^. Importantly, these patterns (Figure 4, Supplementary Figure 2) are not identifiable when analyzed across all mouse models (supplementary figure 5), corroborating the notion that the ASD connectivity landscape is composed of a set of etiology-specific connectivity aberrances converging onto a small set of common networks dysfunction modes. These findings strongly support current efforts of using neuroimaging *per se* ^2^ or as part of multidimensional decompositions ^48^, to deconstruct ASD heterogeneity into homogenous subtypes, and suggests that rsfMRI is sufficiently sensitive to ASD-related pathology to serve as a reliable categorization axis in these efforts.

Interestingly, we found that connectivity aberrations in two of these clusters affect similar networks, but are opposite in direction, i.e. networks that are *over-*connected in one cluster are *under-*connected in the other. As discussed above, this finding may be critical for explaining inconsistencies in human studies that consider all ASD subjects as a single group, without considering the underlying etiological variability. It is indeed conceivable that the combined effect of these two opposite subtypes may reduce -if not cancel out entirely -the ability of rsfMRI to identify significant group-level changes. The exact clinical significance and occurrence of these subtypes remain to be established. However, it is tempting to speculate that such a scenario may partly account for previous reports hinting at the lack of robust connectivity alterations in ASD cohorts ^12,49^.

While the relatively low number of mutations examined here does not permit speculation about the significance of the etiological groups identified in terms of molecular pathways, transductional cascade and common behavioural deficits, it is encouraging to note that some associations may be mechanistically meaningful. These include, for example, the observation that both homozygous and heterozygous CDLK5 mutants were grouped together with MECP2, consistent with the hypothesis that these models share the same molecular dysfunction since CDKL5 is a kinase able to mediate MeCP2 phosphorylation ^50,51^. However, for the most part, the group of mutations clustered together are highly heterogeneous and do not appear to represent any obvious signaling or molecular cascade. This finding is however not surprising, as temporal pleiotropy and developmental timing are essential factors which might cause similar patterns of network dysfunction even if the affected molecular mechanisms are seemingly distinct ^5^. Notably, our clustering approach did not distinguish between mutants with a clear genetic association with ASD versus mutants that code for syndromes which are only partially related to autism. This indicates that the mapping between distinct ASD etiologies and the brain connectivity subtype is complex and most likely driven by a multitude of different biological, environmental and developmental factors. Future expansions of our dataset to a sufficiently large number of genetic etiologies which are more representative of the complex genetic landscape of ASD ^52^ may be envisaged to further explore whether alterations in known molecular pathways converge on similar network alterations. The expansion of the AMC database might not be limited to ASD mutants but could also include mouse models of other neurodevelopmental disorders to identify whether similar alterations of brain connectivity manifest across diagnostic categories. Similarly, the addition of multidimensional phenotyping data to include behavioral readouts may help to assess the translational significance of the mapped brain (dys)connectivity signatures since it might reveal their relationship with core deficits that characterize ASD. In this respect, it is interesting to note that somato-motor, insular and striatal networks appeared to be among the most frequently affected substrates across mutations and clusters. This finding is not surprising in light of the sensory perception and motor-related impairments commonly described in many of the examined models. For example, both Engrailed 2 and Fmr1 knockout mice are characterized by blunt encoding of tactile stimulation frequency, increased sensitivity to somatosensory stimuli and larger size of receptive fields in the somatosensory cortex ^24,53,54^. Similarly, hyperactivity and excessive self-grooming have been described in many models included in this study, most notably Shank3 ^55^ and CNTNAP2 ^56^, and repetitive/restricted behavior has also been observed in En2, Fmr1, BTBR and MECP2 mutants ^57^, recapitulating hallmark ASD symptoms commonly related to compromised fronto-striatal-motor signaling ^58–60^. Future research employing targeted cellular and circuit manipulations in rodents combined with similar fMRI recordings ^61,62^ may crucially uncover the bases of these network-level alterations, their behavioral significance, and their translational relevance with respect to analogous measurement in clinical populations ^11,63^.

In conclusion, our cross-etiological analyses of functional connectivity in 16 ASD models revealed a broad spectrum of functional connectivity aberrancies. Even though there was convergence towards a single signature, several connectivity subtypes were identified. Our results define four prominent network dysfunction modes of cross-etiological relevance for ASD, and highlight a pivotal contribution of etiological variation to connectivity heterogeneity in autism as hinted by previous human studies ^64,65^.

## Materials and methods

### Ethical statement

All study procedures were approved by the institutional review board at the involved medical centers and are in accordance with the ethical standards of the Declaration of Helsinki of 1975, as revised in 2008. All experiments performed at ETH Zürich were in accordance with the Swiss federal guidelines for the use of animals in research, and under licensing from the Zürich Cantonal veterinary office. All experiments performed at IIT Rovereto were conducted in accordance with the Italian Law (DL 27/1992 and DL 26/2014, EU 63/2010, Ministero della Sanità, Roma to A.G.) and the recommendations in the Guide for the Care and Use of Laboratory Animals of the National Institutes of Health. Animal research protocols were also reviewed and consented to by the respective animal care committees.

### Magnetic resonance imaging

Data acquisition was performed in both sites on a Biospec 70/16 small animal MR system (Bruker BioSpin MRI, Ettlingen, Germany). Scans from ETH Zürich were obtained with a cryogenic quadrature surface coil (Bruker BioSpin AG, Fällanden, Switzerland). Scans from IIT Rovereto were obtained with a 72 mm birdcage transmit coil and a custom-built saddle-shaped four-element coil for signal reception. Common standard adjustments included calibration of the reference frequency power and the shim gradients using MapShim (Paravision v6.1). BOLD rsfMRI time series were acquired using an Echo Planar Imaging (EPI) sequence, harmonized between the two sites. Sequence details from different sites/models are described in the appropriate references from table 1. In all datasets, mild anesthesia levels were maintained using either isoflurane+medetomidine or halothane (0.75%) and were based on published protocols optimized for maintaining physiological stability, and include intubation and mechanical ventilation ^19,44,66,67^.

### Data preprocessing and connectome construction

Resting-state fMRI datasets were commonly preprocessed using a standard minimal-preprocessing pipeline ^68^. Briefly, the nuisance model used for signal regression includes six head motion parameters and ventricle signals. Thereafter, datasets were de-spiked, band-pass filtered, skull-stripped and normalized first to an EPI study-specific template and then to the Allen Brain Institute reference atlas (http://mouse.brain-map.org/static/atlas) using ANTs v2.1 (http://picsl.upenn.edu/software/ants/). BOLD time series were extracted using the Allen Reference Atlas ontology and their connectivity couplings were measured using Z-scored regularized Pearson’s correlation coefficient (FSLNets). 165 ROIs in both hemispheres were included in the rsfMRI analysis. This included regions in isocortex, olfactory areas, hippocampal formation, cortical subplate, striatum, pallidum, thalamus, hypothalamus, midbrain and pons (Figure 1A).

### Cluster definition and verification

All statistical analyses and clustering were performed using Matlab (R2019a) and Python (Scikit-learn, UMAP)^69,70^. We first sought to reduce the complexity of the full connectome (165×165 ROIs resulting in 13530 edges) by considering only the strongest, positively-correlated edges. We therefore applied a sparsity threshold to the connectivity matrix. Based on an analysis in non-transgenic mice (see Results), we choose a sparsity of 4% because that ensures high similarity across the different cohorts. This level of sparsity led to the inclusion of 545 connections (i.e. edges). For each of these edges and in each of the 16 ASD mouse models, we quantified deviations from the wildtype’s littermate mean connectivity strength in form of Cohen’s d effect size estimates.

The effect size vectors were then fed into a Gaussian Mixture Model (GMM) clustering algorithm to establish the links between mouse cohorts based on their effect sizes. GMM treats the data as a superimposition of multiple Gaussian distributions and it applies the Expectation-Maximization (EM) algorithm to determine the mean and the variances of these distributions (the latter distinguishes GMM from other clustering algorithms, such as k-means).

To determine the consistency of the clustering, we used a bootstrapping procedure to generate a distribution of the real data which we obtained by applying the GMM after randomly removing 20% of the data in each cohort. We compared this against a *null model*, which was obtained by randomly shuffling the labels of mutant versus WT mice of each cohort. This operation was repeated 1000 times to determine the consistency of the clustering (1000 times). For each iteration, we calculated the effect-size in every edge and applied GMM clustering as described above. The proportion of time (using 1000 repeats) that two mouse cohorts were grouped together was compared between the *bootstrapped* and *null* distributions and used to determine similarity across cohorts in the clusters.

To establish the best cluster solution, we computed the silhouette value. The silhouette value ranges from –1 to 1. A high silhouette value indicates that models are well matched to their own cluster, and poorly matched to other clusters. We measured silhouette scores for different numbers of clusters (repeated 1000 times), and compared the values between *bootstrapped* and *null* distributions.

To assess connectivity profiles on a mid-level scale, the number of edges that turned significant (randomized, nonparametric statistics, p<0.05) in/out of a given anatomic structure were summed (node-level comparison). Finally, for the coarsest-scale analysis, anatomic regions were collapsed (summed) into their parent structures according to the Allen Mouse Brain Common Coordinate Framework (CCFv3) ontology (parent-level comparison).

For statistical analysis of connectome edge-strength distributions, we used a non-parametric permutation test using FSL randomize (n=5000 repeats) corrected for multiple comparisons using false detection rate (FDR).

---

**Supplementary Figure 1.**
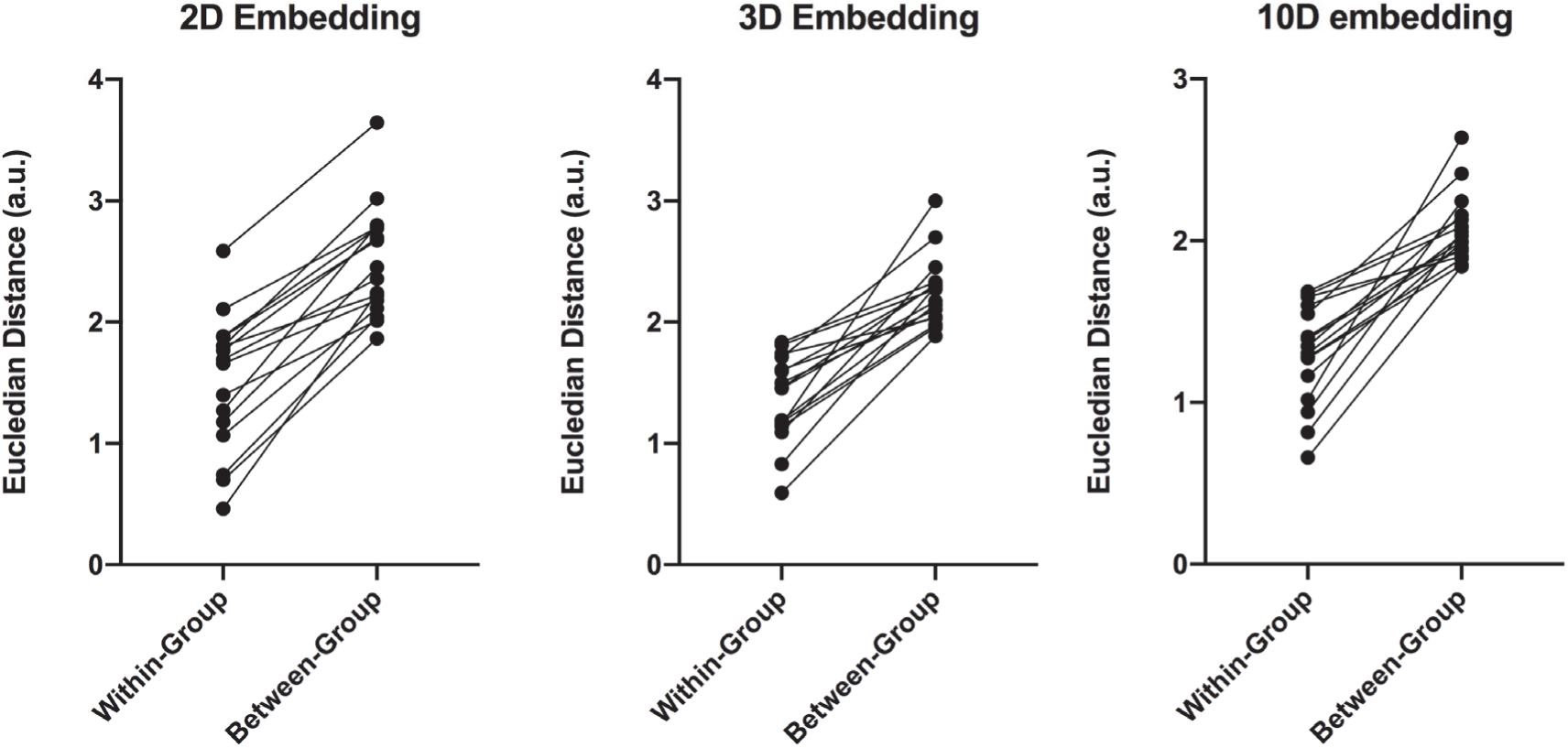
Averaged Euclidian distance within-and between-groups in the UMAP two-dimensional, three-dimensional and ten-dimensional embeddings. In all conditions, mouse models show a reduced distance between animals of the same cohort compared to animals of other cohorts, suggesting that similar connectivity deviations depend on the etiology of the model.

**Supplementary figure 2.**
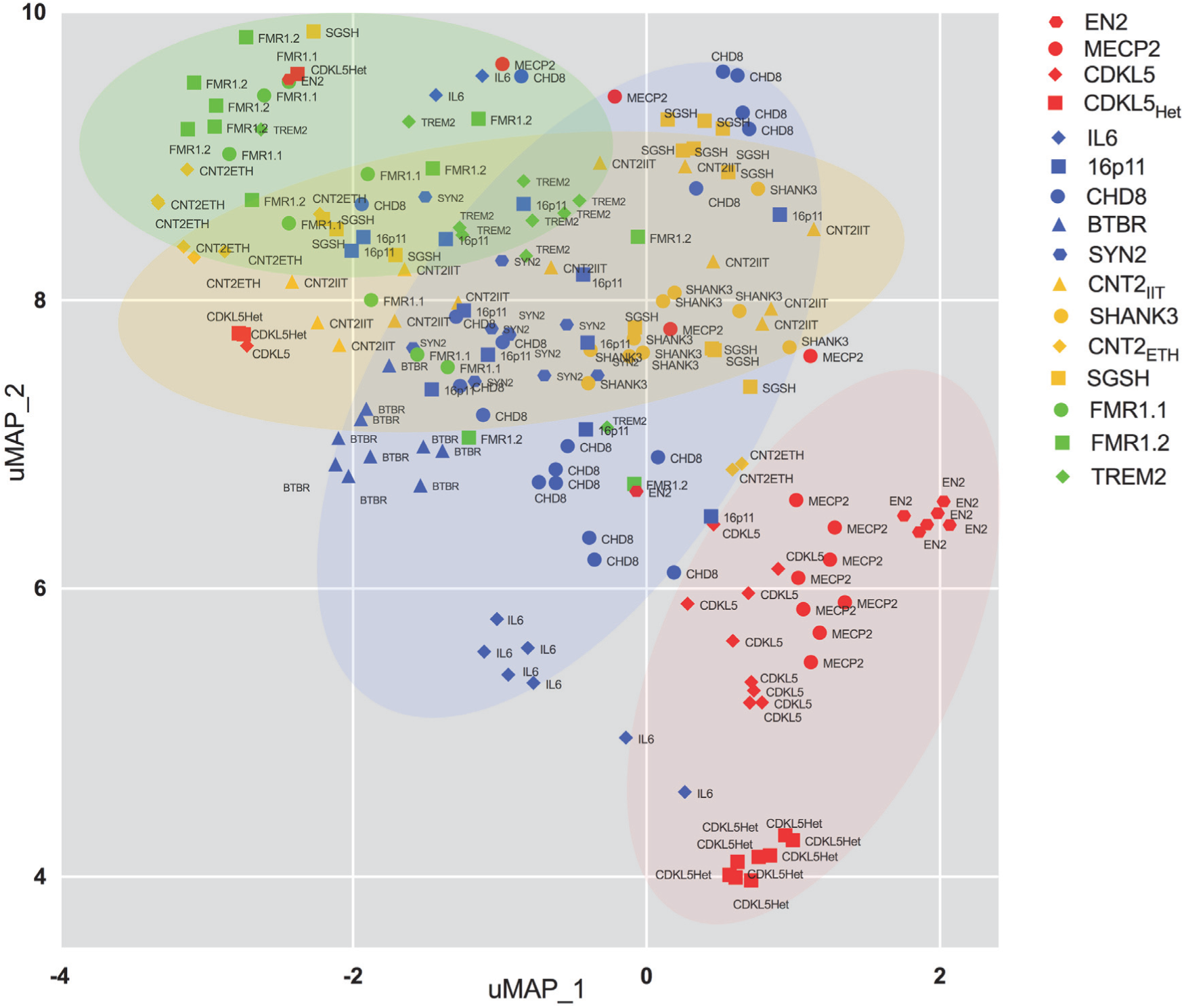
UMAP 2-dimensional embedding of the connectome data from the 176 individual animals of the AMC dataset. Individual data are Z-scored and normalized to the average cohort’s wildtype control population. The shape of the elements represents the cohort, while the colors represent the clusters.

**Supplementary figure 3:**
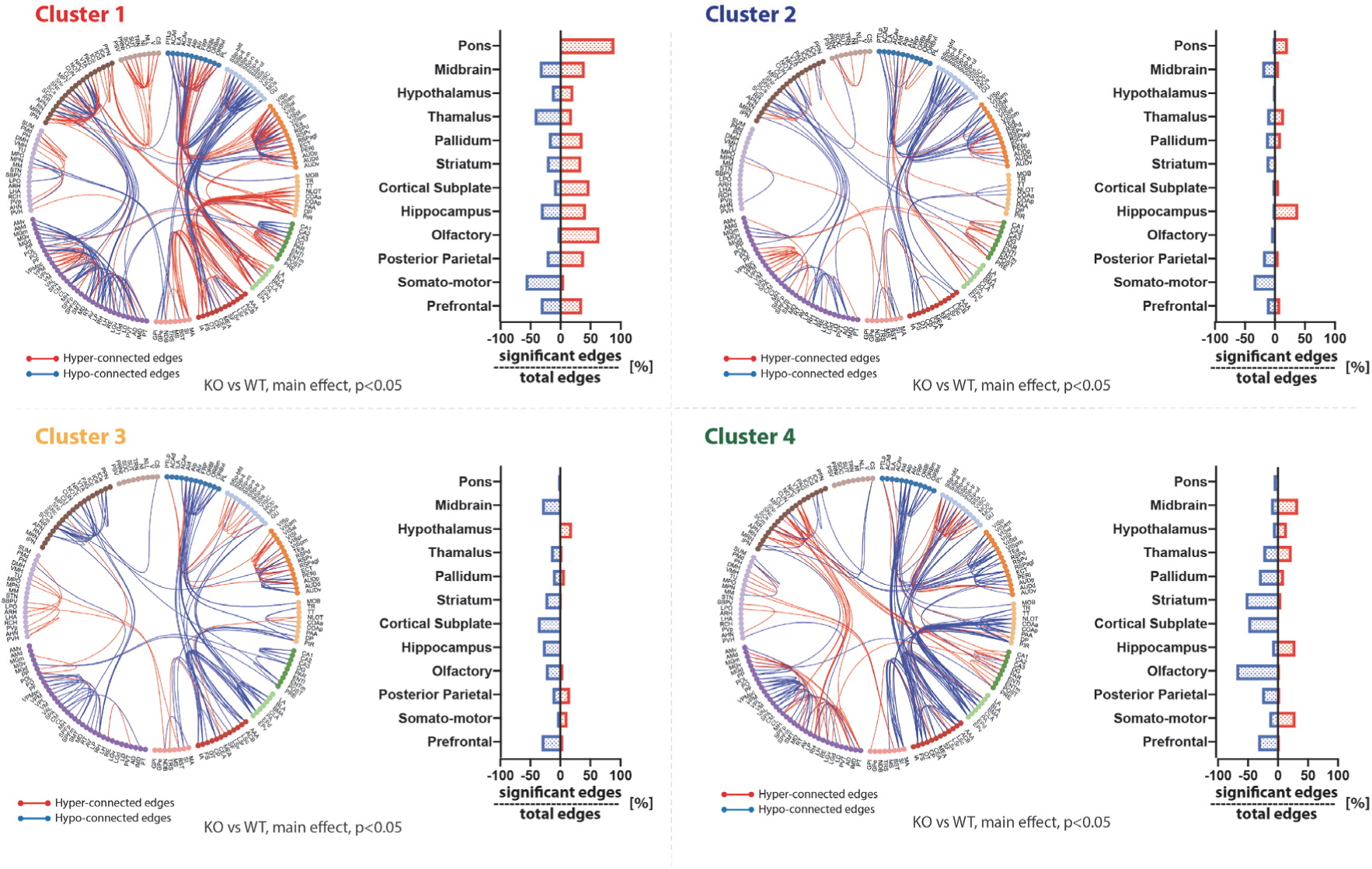
edge-edge and parent-level statistics show a brain-wide heterogeneous distribution of under-and over-connectivity across the four clusters.

**Supplementary Figure 4.**
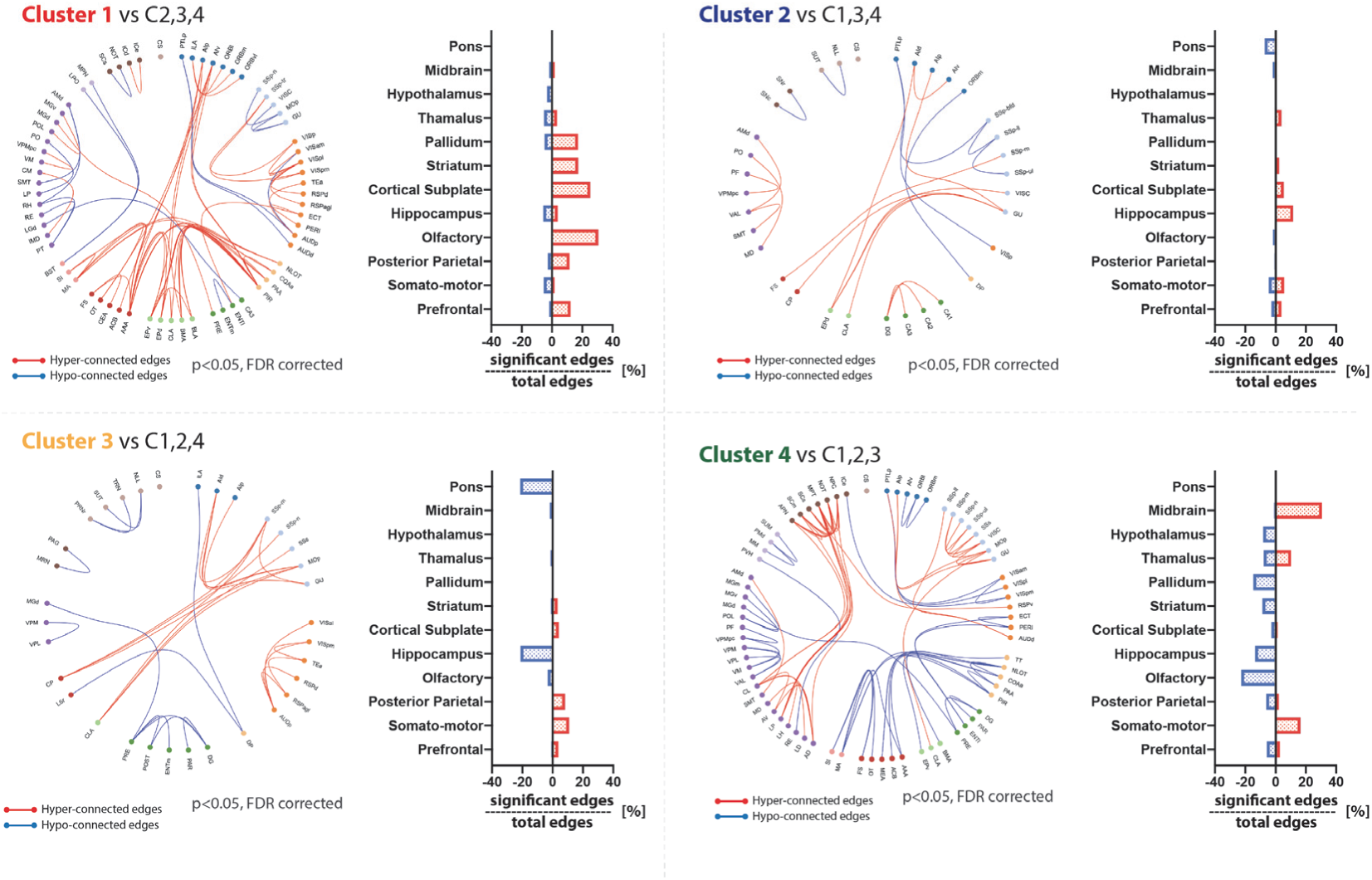
Edge-edge and parent-level statistics indicate under-and over-connectivity deficits that are unique to one cluster as compared to the other three. This analysis highlights the unique profile of each cluster. The ASD-etiologies in Cluster 1 are characterized by prominent over-connectivity in the prefrontal cortex, posterior parietal, olfactory area, cortical subplate, striatum and pallidum. This was in complete opposition to the alterations observed in Cluster 4, which are characterized instead by under-connectivity in the same areas, but also by a strong over-connectivity within the somatomotor areas and between midbrain and thalamus. Cluster 2 and 3 instead showed increased connectivity in hippocampus and decreased connectivity in pons, or a marked reduction in connectivity in pons and hippocampus, respectively.

**Supplementary figure 5.**
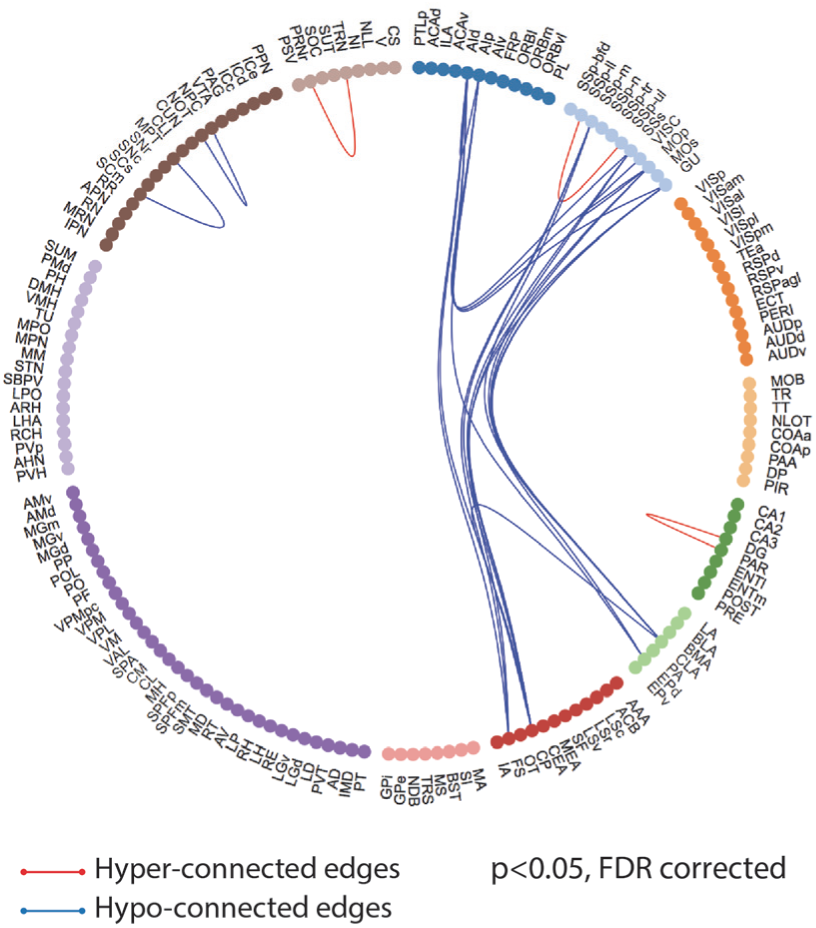
Non-parametric statistics revealed a handful of connectivity deficits that are common across all models. Circos plot illustrates that the majority of the significant edges have under-connectivity between insula, somatosensory cortex, stratum, amygdala. Blue colors represent under-connectivity. Red colors represent over-connectivity as compared to wildtype littermates. AIp = Anterior Insula, posterior, Cla=claustrum, CPu = caudoputamen, DG = dentate gyrus, EPd= Endopiriform nucleus, dorsal part, GU=gustatory area, MOp=primary motor area, PAR= parasubiculum, SCm= superior colliculus, motor related, SSp-ul= primary somatosensory area, upper limb, SSs = supplementary somatosensory area.

